# Investigation on key aspects of mating biology in the mosquito *Aedes koreicus*

**DOI:** 10.1101/2022.12.01.518615

**Authors:** Silvia Ciocchetta, Francesca D Frentiu, Fabrizio Montarsi, Gioia Capelli, Gregor J Devine

## Abstract

*Aedes koreicus* is a mosquito (Diptera: Culicidae) from Northeast Asia with a rapidly expanding presence outside its original native range. Over the years the species has been discovered in several new countries, either spreading after first introduction or remaining localised to limited areas. Notably, recent studies have demonstrated the ability of the species to transmit zoonotic parasites and viruses both in the field and in laboratory settings. Combined with its invasive potential, the possible role of *Ae. koreicus* in pathogen transmission highlights the public health risks of its invasion. In this study, we used a recently established population from Italy to investigate aspects of biology that influence reproductive success in *Ae. koreicus*: autogeny, mating behaviour, mating disruption by the sympatric invasive species *Aedes albopictus*, and the presence of the endosymbiont *Wolbachia pipientis*.

Our laboratory population did not exhibit autogenic behaviour and required a blood meal to complete its ovarian cycle. When we exposed *Ae. koreicus* females to males of *Ae. albopictus*, we observed repeated attempts at insemination and an aggressive, disruptive mating behaviour initiated by males. Despite this, no sperm was identified in *Ae. koreicus* spermathecae*. Wolbachia* was not detected in this *Ae. koreicus* population and therefore had no effect on *Ae. koreicus* reproduction.

## Introduction

After its first detection in Belgium in 2008 (Versteirt *et al*., 2012), the mosquito *Aedes koreicus*, commonly known as the invasive Korean bush mosquito, has invaded and established in several states in Europe and European neighbouring countries (ECDC, 2022). In some areas, such as Italy, the species is currently expanding its distribution (Gradoni *et al*., 2021; Negri *et al*., 2021; Arnoldi *et al*., 2022), but in others, such as Germany, it has shown a relatively low tendency to spread despite suspected repeated introductions (Hohmeister *et al*., 2021; Kurucz *et al*., 2022).

While the role of *Ae. koreicus* in arthropod-borne diseases transmission is still largely unclear, the species is known to vector dog heartworm *Dirofilaria immitis* (Filarioidea: Onchocercidae) under laboratory conditions (Feng, 1930; Montarsi *et al*., 2014), a finding later supported by field evidence of filarial DNA in *Ae. koreicus* sampled near the city of Pécs (Baranya County) in Hungary (Kurucz *et al*., 2018). *Ae. koreicus* infection with *Wuchereria bancrofti* (Filarioidea: Onchocercidae) has also been documented (Yamada, 1927), and this mosquito may have a role as an intermediate host for *Brugia malayi* (Filarioidea: Onchocercidae) to infect humans (KCDC, 2007). The potential of *A. koreicus* to transmit chikungunya virus was demonstrated for the first time under laboratory conditions by Ciocchetta *et al*. (Ciocchetta et al., 2018). This study showed how virus transmission was temperature-dependent, and results were further confirmed by Jansen *et al*. (Jansen *et al*., 2021). The same study reported a low vector competence for Zika virus and no transmission of West Nile virus. A few studies have mentioned *Ae. koreicus*’ ability to transmit Japanese encephalitis virus (JEV) in the laboratory and in the field (Miles, 1964; Gutsevich *et al*., 1971; Takashima & Rosen, 1989). However, JEV was not detected in *Ae. koreicus* collected in Korea during more recent monitoring activities (Gutsevich *et al*., 1970; Kim *et al*., 2005; Kim *et al*., 2007).

Even though *Ae. koreicus* was first detected in Europe more than 14 years ago, its mating biology remains largely unknown. Reproductive success plays a fundamental role in mosquito establishment and population growth (Clements, 1992; Juliano & Lounibos, 2005; Takken *et al*., 2006) and an assessment of the reproductive biology of *Ae. koreicus* could assist in determining its invasive potential. In this study, we investigated several important aspects that may influence the mating biology and reproductive success of the Korean bush mosquito in Italy, such as autogeny, mating behaviour and competitive mating with a sympatric invasive mosquito species (*Aedes albopictus*). We also screened the mosquito population used to derive our colony for the presence of the endosymbiont *Wolbachia pipientis*.

In some hematophagous arthropods, such as mosquitoes, completion of an ovarian cycle and the production of viable offspring can occur in the absence of a blood meal in a process called autogeny (Roubaud, 1929), most likely as a survival strategy when hosts are rare (Lucius *et al*., 2017). Autogeny is hypothesised to allow the persistence of a population when the presence of vertebrate hosts is low, or to allow for rapid growth of a mosquito population at the start of a season (O’Meara, 1985; Reisen & Milby, 1987). This allows mosquitoes to persist in uncertain environments and rapidly exploit optimal conditions; however, the number of eggs laid might vary considerably compared to eggs laid after a blood meal (O’Meara & Krasnick, 1970; O’Meara & Edman, 1975; Mulla, 1997). Furthermore, this behaviour may delay contact with infected hosts, and could therefore impact transmission of human pathogenic viruses by mosquito vectors early in the season. Autogeny may be facultative or obligate depending on the species and environmental conditions (O’Meara & Krasnick, 1970; O’Meara & Edman, 1975).

The autogeny phenotype has been demonstrated both in the Culicinae and Anophelinae mosquitoes (Clements, 2013). Within the Culicinae group, autogeny is commonly reported in the genus *Culex* (Provost-Javier *et al*., 2010), and limited levels of autogeny have been observed in numerous species of *Aedes* mosquitoes (Rioux *et al*., 1975), including some of the main mosquito threats of this century, *Ae. albopictus* and *Aedes aegypti* (Trpis, 1977; Chambers & Klowden, 1994; Mori *et al*., 2008; Gulia-Nuss *et al*., 2015; Aardema & Zimmerman, 2021). An essential component of autogeny is the female mating status (evidence that sperm transfer occurred): egg development in certain mosquito species does not initiate unless mating occurs, and male accessory gland products can play a central role for oogenesis (O’Meara & Evans, 1976, 1977). The ability to identify sperm in the *Ae. koreicus* female reproductive tract (mating status) is necessary to identify whether the absence of autogeny is simply the result of non-mated females. It is also fundamental in evaluating mating behaviour, reproductive success, and the subsequent spread of invasive species in a new territory.

The establishment of an exotic species may be hampered by the disruption of conspecific mating by the aggressive mating behaviour of males of different species (Tripet *et al*., 2011) and by interspecific cross-insemination (satyrization) (Lounibos, 2007; Alto & Lounibos, 2013). Satyrization (Ribeiro & Spielman, 1986) is a form of sterility caused by interspecific mating. For example, the transfer of *Ae. albopictus* male accessory gland product to *Aedes aegypti* females causes them to become refractory to further mating (including with conspecific males) (Nazni *et al*., 2009; Tripet *et al*., 2011; Lima-Camara *et al*., 2013). Although *Ae. albopictus* males are particularly efficient in satyrizing *Ae. aegypti* females, similar interactions have been noted between *Ae. albopictus* and other *Aedes* species such as *Aedes polynesiensis* and members of the *Aedes scutellaris* group (Gubler, 1970; Ali & Rozeboom, 1971a, 1971b).

Additionally, mosquito reproductive behaviour can be influenced by the presence of the endosymbiotic bacteria *Wolbachia pipientis. Wolbachia* are small (0.5–1μm), intracellular, α-proteobacteria originally identified from the ovaries of *Culex* mosquitoes in 1924 (Hertig & Wolbach, 1924) and known to infect the reproductive organs of 40-60% of insect species (Jeyaprakash & Hoy, 2000; Hilgenboecker *et al*., 2008; De Oliveira *et al*., 2015; Weinert *et al*., 2015). They can affect host reproduction by increasing the reproductive success of infected females, thus enhancing the bacteria’s maternal transmission and changing male sperm structure such that only mating with a male infected by the same bacterial strain will lead to progeny (a mechanism called cytoplasmic incompatibility) (Werren *et al*., 2008). In some cases *Wolbachia* can induce parthenogenesis (Stouthamer et al., 1999), and influence fecundity (Alexandrov *et al*., 2007) and oogenesis (Dedeine *et al*., 2001; Dedeine *et al*., 2003). Our aim here was to provide the basis for further studies on the reproductive behaviour of *Ae. koreicus* and its potential to become established when introduced in new territories.

## Methods

### Determination of autogeny in Aedes koreicus

*Aedes koreicus* larvae were obtained from a colony maintained at the QIMR Berghofer Medical Research Institute (QIMRB) (Ciocchetta *et al*., 2017). Eggs laid on Masonite^®^ sticks were hatched in rainwater. Due to the low hatching rate of this species (Ciocchetta *et al*., 2017), larvae were obtained from colony eggs pooled in order to produce sufficient adults for experimentation. Pupae developed from larvae after nine days and were sexed using the method previously described (Ciocchetta *et al*., 2017). To generate three experimental replicates, male and female pupae were placed together in three different cages (BugDorm^®^ Insect Rearing Cage, 30 x 30 x 30 cm) at the following initial numbers: cage 1, 161 males - 163 females; cage 2, 161 males - 174 females; cage 3, 161 males - 170 females.

The cages of adults were maintained in environmental chambers (Panasonic, Osaka, Japan) as described previously (Ciocchetta *et al*., 2017). A 10% w/v sucrose solution was provided *ad libitum* and each cage was equipped with one egg collection tray (© 2014 Genfac Plastics Pty Ltd, 18.3 x 15.2 x 6.5 cm) with rainwater and Masonite^®^ sticks as oviposition substrates (Figure 1). The position of the cages within the environmental chamber was changed twice per week to minimise positional bias. The number of emerging adults was counted, and cages were checked daily for eggs. After three weeks of caging, one of the three cages was randomly chosen (cage 2) to proceed to blood feeding on human volunteers (QIMRB Human Research Ethics Committee approval HREC361). The percentage of fed mosquitoes was recorded. Two weeks after blood feeding (and seven days from the start of oviposition), eggs were collected, counted, and stored in an anti-leak plastic bag. Additionally, 5 female mosquitoes from the blood-fed cage and 10 female mosquitoes from the remaining two cages were killed (using CO_2_), and their ovaries were dissected in a drop of phosphate-buffered saline (PBS) on a glass slide at a magnification of 10x in order to identify mature follicles (stage IVb and V) (Christophers, 1911; Clements & Boocock, 1984; Armbruster & Hutchinson, 2002; Hugo *et al*., 2003; Itina *et al*., 2014). The viability of a subsample of eggs collected from cage 2 (n= 1189) was measured after 14 days of storage (Ciocchetta *et al*., 2017) to verify the successful completion of the gonotrophic cycle in that cage. Observation of Masonite^®^ sticks for presence of eggs in the non-blood-fed cages continued until all adult mosquitoes had died and the absence of autogeny was confirmed.

**Figure 1.**
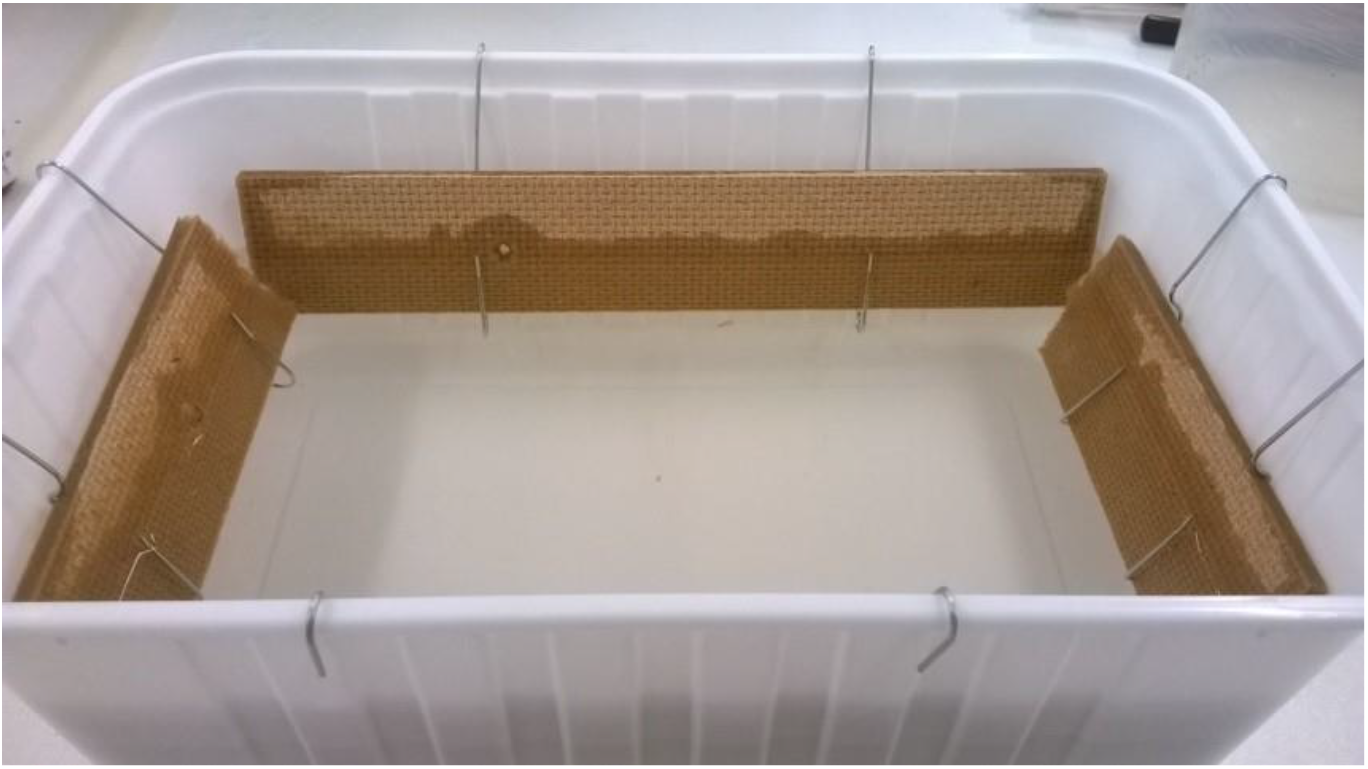
Egg collection tray with rainwater and Masonite^®^ sticks.

### Conspecific Aedes koreicus mating behaviour

*Aedes koreicus* pupae were derived from mosquito eggs laid on Masonite^®^ sticks and sexed according to Ciocchetta et al. (Ciocchetta *et al*., 2017). 190 males and 240 females were separated into two different BugDorm^®^ cages placed in environmental chambers for emergence, at the previously described colony rearing temperature and relative humidity (Ciocchetta *et al*., 2017). Preliminary observations demonstrated that *Ae. koreicus* mosquitoes mate under conditions of scarce illumination (Silvia Ciocchetta, personal observation). As a result, the light/dark cycle was reversed so that mosquito behaviour could be observed under crepuscular and dark conditions. The observation cage was a modified BugDorm^®^ cage with transparent plexiglass used on one side of the cage instead of mesh. Male mosquitoes require a sufficient period of time for genitalia and sperm development before mating (Oliva *et al*., 2014b), whereas females are often receptive as soon as they emerge (Takken *et al*., 2006). As a result, 6-7-day old virgin males and 2-3-day old virgin females were caged together, and their behaviour recorded. At 12-13 hours intervals, 25 females were aspirated from the experiment cage, anaesthetised with CO_2_ and dissected in a drop of phosphate-buffered saline (PBS) on a glass slide at 10x magnification. A cover slip was used to rupture the spermathecae and allow for sperm visualisation at an increased magnification of 40x.

### Preliminary observations of Aedes albopictus and Aedes koreicus mating disruption

*Aedes koreicus* larvae were reared as previously described (Ciocchetta *et al*., 2017). *Aedes albopictus* larvae (from a colony established at QIMR from eggs collected on Hammond Island, Torres Strait, Australia, in May 2014) were similarly reared, but at a temperature of 27 ± 1°C. The colonies of both species were synchronised to pupate at the same time. Pupae were individually placed in Falcon^®^ tubes containing 5 to 10 ml of rainwater to allow the collection of emerging virgin males or females. 3-4 days old *Ae. albopictus* males (N=27) ready for copula (Oliva *et al*., 2014b) and 2-3 days old virgin *Ae. koreicus* females (N=22) were introduced in a BugDorm^®^ cage containing a solution of 10% w/v sucrose. The interaction between the two mosquito species was recorded utilising a GoPro^®^ Hero 3 camera. After five days, all female mosquitoes were anesthetised with CO_2_ and the spermathecae were dissected in a drop of saline buffer, crushed under a cover slip and scanned at 40x magnification for the presence of sperm.

### Wolbachia presence in field-collected Aedes koreicus

Field-collected *Aedes koreicus* sampled during a survey carried out in north-eastern Italy from 2011 to 2015 (Montarsi *et al*., 2015), form the same population used to derive our QIMRB colony (Ciocchetta *et al*., 2017), were screened for the presence of *Wolbachia pipientis*. Females (n=21) collected in Belluno (46°08’44.3”N 12°12’38.0”E) in July 2014, were preserved in RNALater^®^ (Invitrogen™), and stored at −80°C. DNA was extracted using QIAGEN DNeasy^®^ Blood and Tissue Kit. The extracted DNA was utilised as a template for the polymerase chain reaction (PCR) targeting the *Wolbachia-specific wsp* and 16s genes and the mosquito housekeeping *RpS17* gene, which acted as a positive control for the extraction: (*wsp* F: 5’– TGGTCCAATAAGTGATGAAGAAAC–3’, R: 5’– AAAAATTAAACGCTACTCCA–3’; *16s* F: 5’–TTGTAGCCTGCTATGGTATAACT–3’, R: 5’– GAATAGGTATGATTTTCATGT–3’; *RpS 17* F: 5’–TCCGTGGTATCTCCATCAAGCT–3’, R: 5’–CACTTCCGGCACGTAGTTGTC–3’) (O’Neill *et al*., 1992; Braig *et al*., 1998; Cook *et al*., 2006).

PCR with *wps* primers was performed using a Phusion^®^ High-Fidelity PCR Kit with initial denaturation at 98°C for 30 sec, followed by a 34 cycles consisting of 98°C for 10 seconds, 59°C for 30 seconds, and 72°C for 30 seconds and a final extension step at 72°C for 10 minutes. The same protocol was applied with *16s* and *RpS17* primers, but the annealing temperatures were 56°C for *16s* primers and 58°C for *RpS17* primers.

DNA for four *Wolbachia*-positive controls was extracted from *w*Mel-infected *A. aegypti* maintained in the QIMR Berghofer insectary (Ulrich *et al*., 2016) using the same extraction kit of the target samples. In each PCR, a sample from an *Ae. aegypti* wildtype colony (QIMRB) that was negative for *Wolbachia* was also tested. DNA from *Culex sitiens* mosquitoes (n=3) infected with *Wolbachia* (QIMRB colony) was extracted using QuickExtract™ DNA Extraction Solution (Epicentre Technologies Corporation) and tested as an additional positive control.

## Results

### Determination of autogeny in Aedes koreicus

Proportion of male: female totals were 123:134, and 103:138 in the two non-blood fed cages (cages 1 and 3), and 116:146 in the 3^rd^, blood fed cage (cage 2). No eggs were observed in the oviposition trays of the three cages for 21 days after co-caging. After this period, a volunteer fed the mosquitoes in the cage designated to be blood fed (97.2% fed, n= 109) and oviposition on the Masonite^®^ sticks in that cage occurred seven days post-feeding. Mature follicles were observed in all 5 mosquitoes dissected from that cage. No eggs were observed on Masonite^®^ sticks in cages 1 and 3, and no mature follicles were found in the female mosquitoes dissected from those cages. The percentage of male and female mosquitoes still alive at the time of these observations (28 days after co-caging) were: cage 1= males 8.9% (n = 123), females 59.7% (n = 134); cage 2 = males 44.8% (n = 116), females 67.1% (n = 146); cage 3 = males 25.2% (n = 103), females 71.0% (n = 138). A total of 4,925 eggs were counted under the stereoscope from the Masonite^®^ sticks collected from the blood fed cage; the average eggs/female = 50.25 was consistent with a previously reported fecundity index (Ciocchetta *et al*., 2017).

### Conspecific Aedes koreicus mating behaviour

No sperm was observed in spermathecae from female mosquitoes dissected 12 ± 0.5 and 25 ± 0.5 hours after co-caging. Mating activity was observed after 25.5 hours showing *Aedes koreicus* males and females in the act of copula and documented with a smartphone device (Oppo F1 Android smartphone; supplemental file 1: Ae. koreicus mating.mp4). Evidence of motile sperm in *Ae. koreicus* female spermathecae (Figure 2) was found in 28% of females (n=25) sampled 31 hours after co-caging with males (females were sampled approximately five hours after evidence of mating activity in the cage to allow the sperm a sufficient period to reach the spermathecae (Oliva *et al*., 2014b).

**Figure 2.**
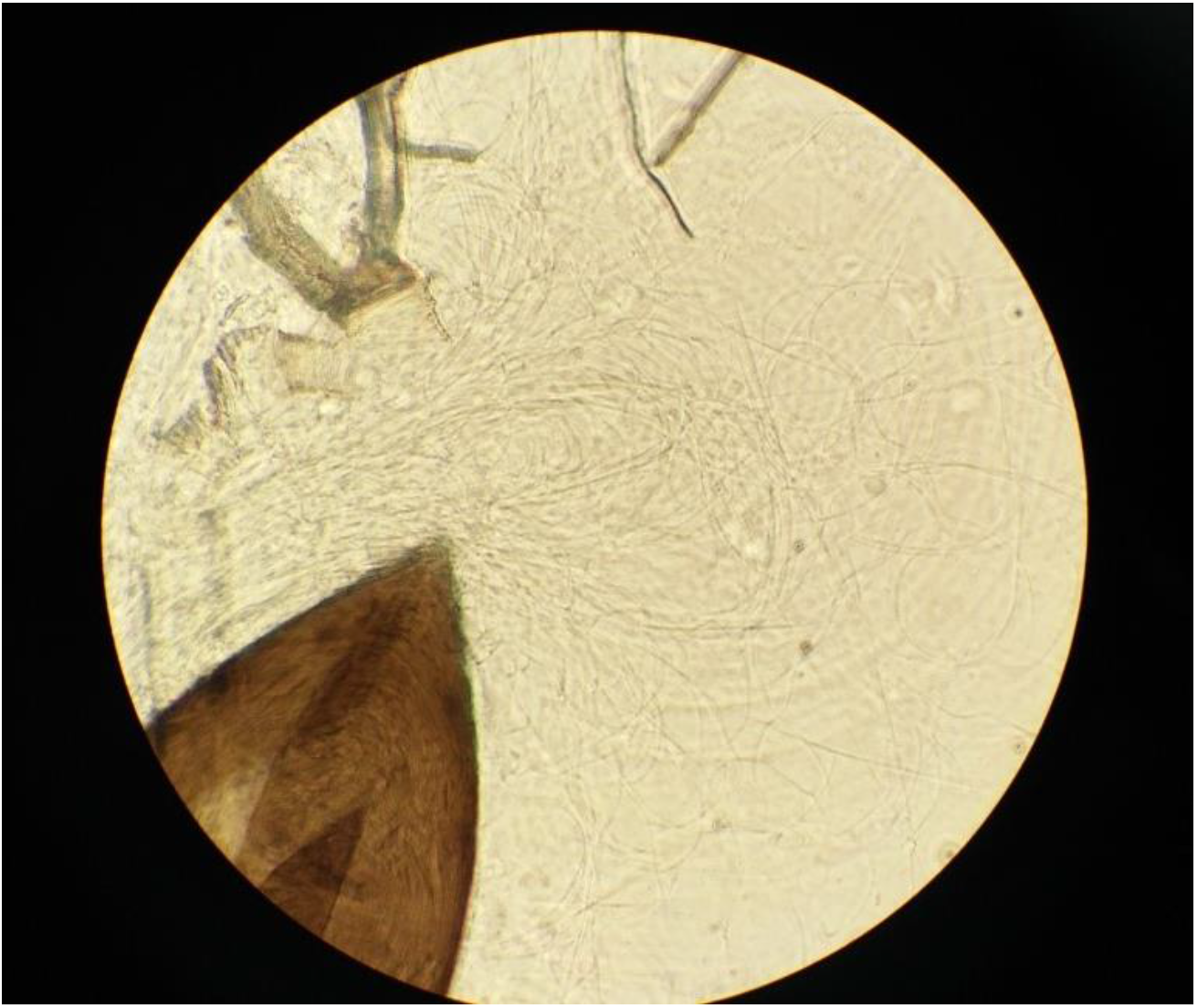
*Ae. koreicus* sperm visible after spermathecae rupture.

### Preliminary observations on Aedes albopictus and Aedes koreicus mating disruption

Despite repeated interactions between *Aedes koreicus* females and *Aedes albopictus* males (supplemental file 2: Ae. albopictus_Ae.koreicus interaction.mp4), no sperm was detected in the 22 individuals dissected (Figure 3). Differences in the size of the mosquitoes (*Ae. koreicus* females were visibly bigger than *Ae. albopictus* males; Figure 4) may have been a possible cause for the failure in interspecific insemination. Although the specific size of each individual was not measured, *Ae. koreicus* females wing length for individuals reared according to previous work (Ciocchetta et al., 2017) has been reported to be over 3 mm; in their work Pudar *et al*. (Pudar *et al*., 2021) reported an *Ae. albopictus* male wing length of approximately 2mm when the species was reared at a temperature close to our experiment (28 ± 1 °C). When reared at the same rearing conditions, these two species maintained their differences in dimensions, with *Ae. koreicus* bigger in size when compared to *Ae. albopictus* (Baldacchino et al., 2017), supporting our observations.

**Figure 3.**
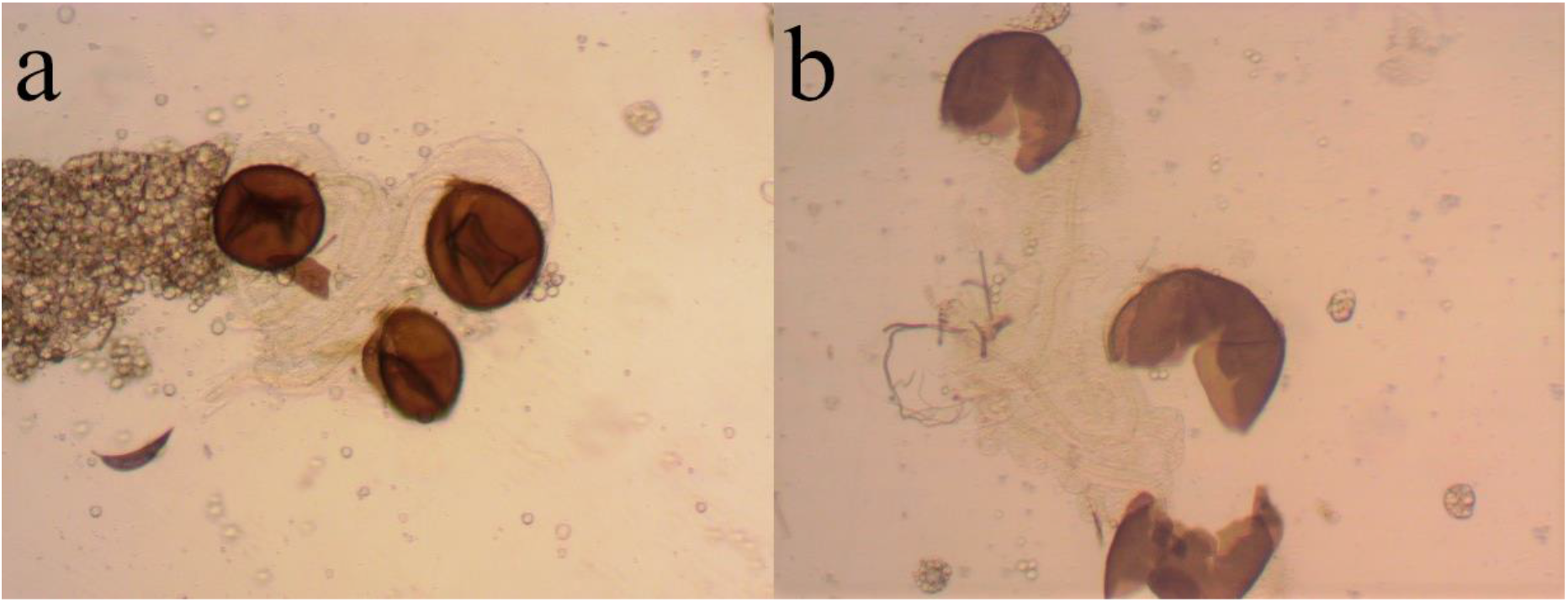
No evidence of *Ae. albopictus* sperm in *Ae. koreicus* spermathecae (a) before and (b) after rupture.

**Figure 4.**
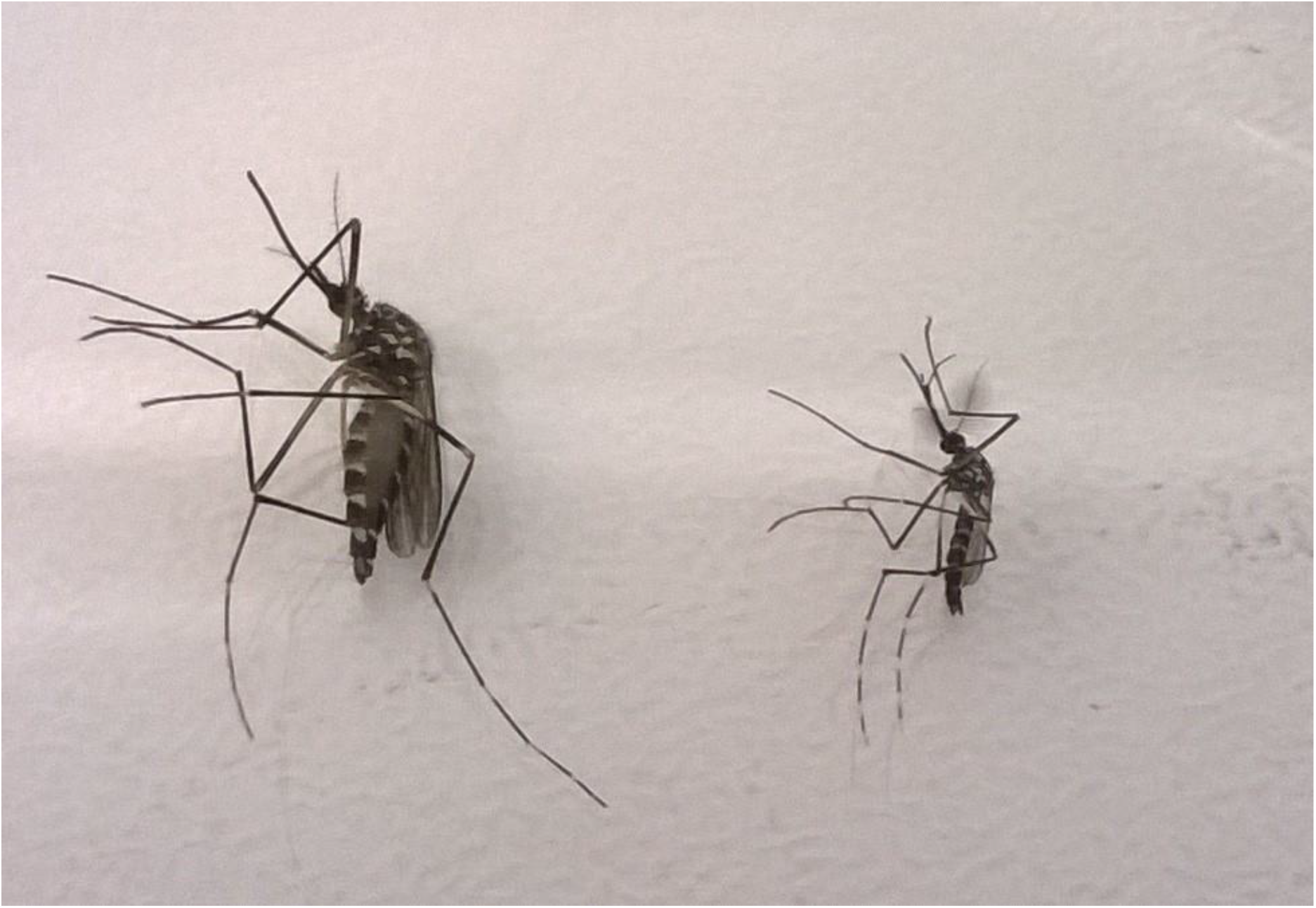
Difference in size between *Ae. koreicus* female (left) and *Ae. albopictus* male (right).

### Wolbachia presence in field-collected Ae. koreicus

No *Wolbachia* was identified in the *Aedes koreicus* field samples. The DNA extraction was validated by running a PCR analysis using *RpS17* housekeeping gene primers for mosquito DNA (Figure 5–7), and *Wolbachia* was detected by the *wsp* and *16S* primers in all positive controls.

**Figure 5.**
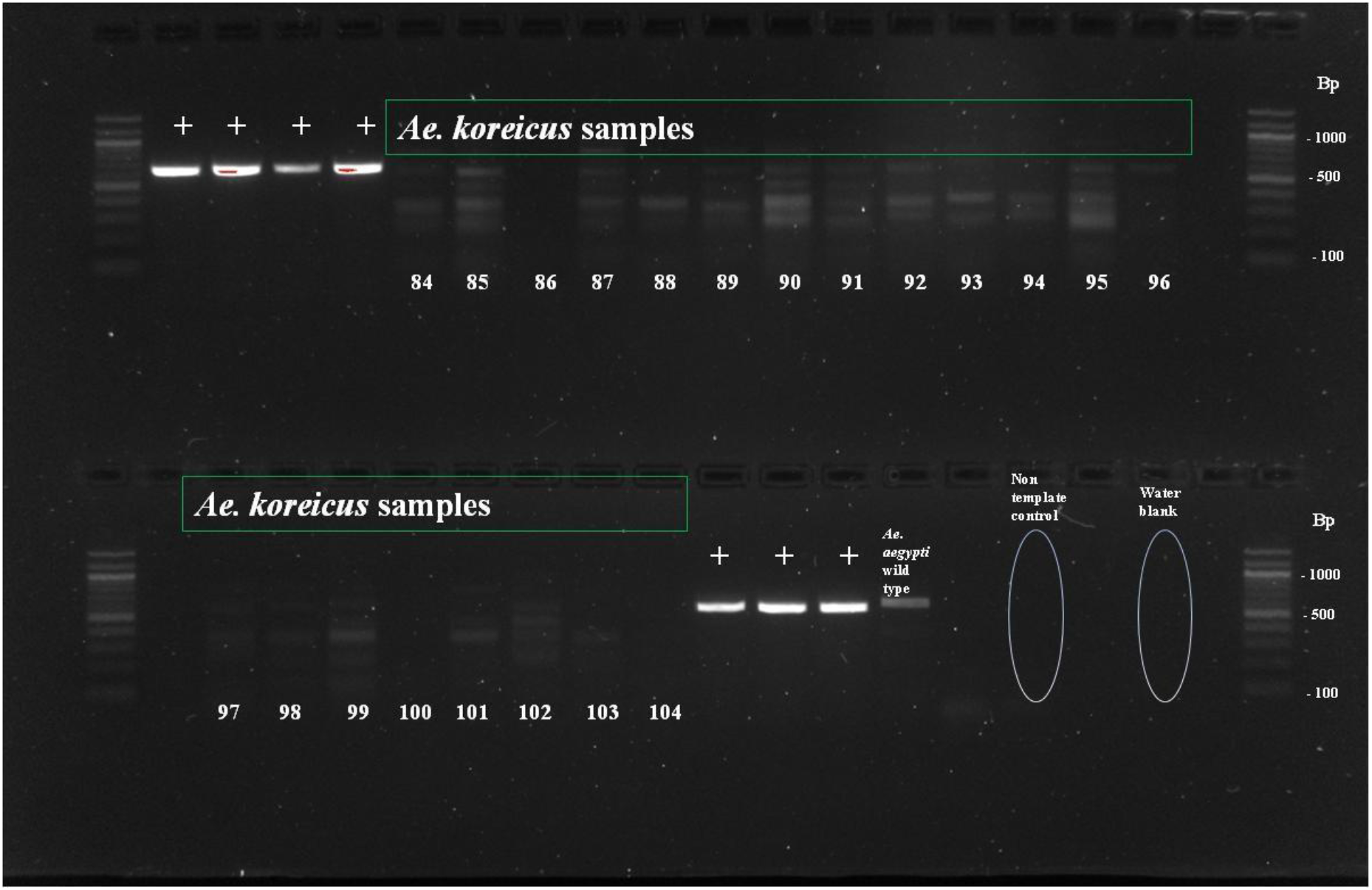
PCR products produced by amplifying *Ae. koreicus* DNA with oligonucleotide primers corresponding to *Wolbachia* gene *wsp*. (a) positive controls indicated by the symbol + (*Ae. aegypti Wolbachia* infected, QIMR Berghofer), *Ae. koreicus* samples lanes 84 to 96; (b) *Ae. koreicus* lanes 97 to 104, positive controls indicated by the symbol + (*Culex sitiens Wolbachia* infected, Chen Wu, QIMR Berghofer), *Ae. aegypti* wildtype *Wolbachia-free*, QIMR Berghofer.

**Figure 6.**
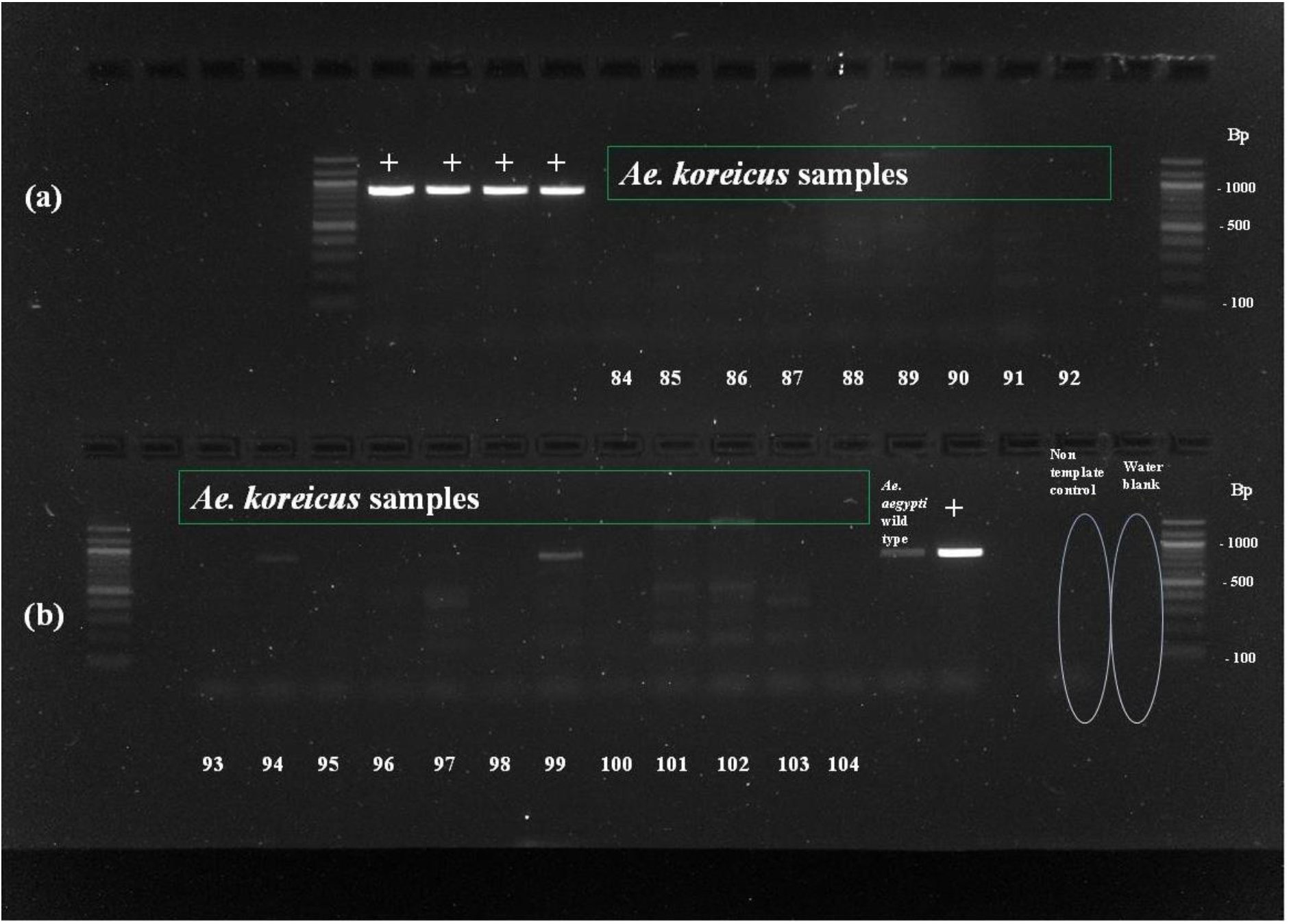
PCR products produced by amplifying *Ae. koreicus* DNA with oligonucleotide primers corresponding to *Wolbachia* gene *16S*. (a) positive controls indicated by the symbol + (*Ae. aegypti Wolbachia* infected, QIMRB), *Ae. koreicus* lanes 84 to 92; (b) *Ae. koreicus* lanes 93 to 104, positive controls indicated by the symbol + (*Culex sitiens Wolbachia* infected, Chen Wu, QIMRB), *Ae. aegypti* wildtype *Wolbachia-free*, QIMRB.

**Figure 7.**
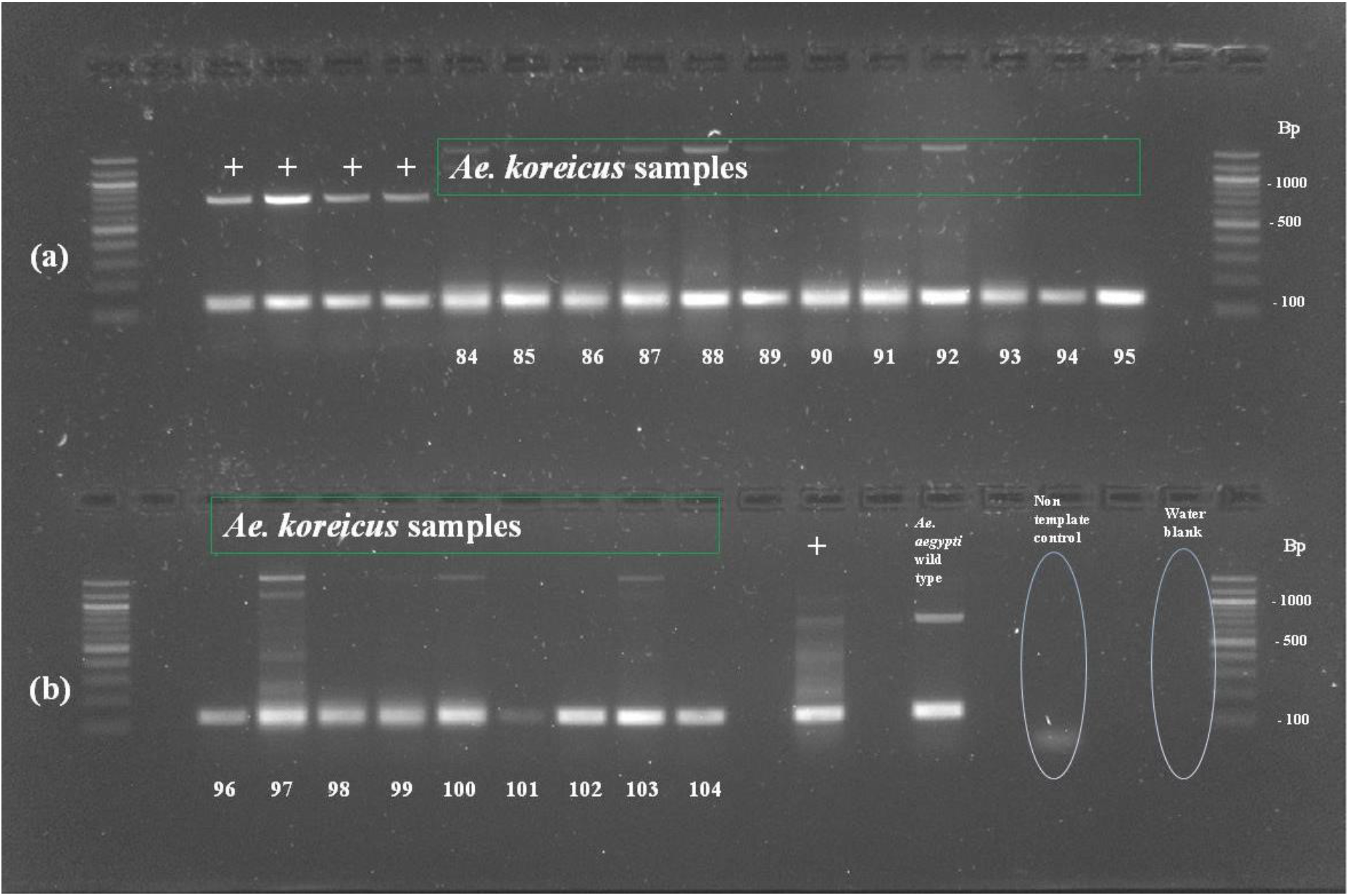
PCR products produced by amplifying *Ae. koreicus* DNA with oligonucleotide primers corresponding to the housekeeping gene *RsP17*. (a) positive controls indicated by the symbol + (*Ae. aegypti Wolbachia* infected, QIMR Berghofer), *Ae. koreicus* lanes 84 to 95; (b) *Ae. koreicus* lanes 96 to 104, positive controls indicated by the symbol + (*Culex sitiens Wolbachia* infected, Chen Wu, QIMR Berghofer), *Ae. aegypti* wildtype *Wolbachia*-free, QIMR Berghofer.

## Discussion

Defined as the ability to produce offspring in the absence of a blood meal, autogeny can influence the vector potential of a mosquito by affecting the abundance or persistence of vectors, even in the absence of immediate hosts (Reisen & Milby, 1987; Tsuji *et al*., 1990). Conversely, autogeny may limit contact with hosts and reduce transmission risks (Spadoni *et al*., 1974; Reisen & Milby, 1987). Our results suggest that *Ae. koreicus* mosquitoes do not display this phenotype under the conditions of our experiment. There was no oviposition when mosquitoes were deprived of a blood source. In early studies with the mosquito *Aedes taeniorhynchus*, O’Meara *et al*. (O’Meara & Evans, 1976) showed that mating may increase the levels of autogeny and that the expression of autogeny is correlated to the environmental conditions in which the larval stages develop and the geographical origin of the population (O’Meara, 1979). In *Ae. taeniorhynchus*, mating was necessary only when larvae were exposed to conditions unfavourable to their development and was otherwise not required for the production of viable eggs (O’Meara, 1979). The observation of *Ae. koreicus* mating behaviour and the detection of sperm in *Ae. koreicus* spermathecae confirmed that the absence of autogeny was not due to a lack of mosquito mating. Moreover, autogenic populations of *Ae. japonicus*, a species phylogenetically close to *Ae. koreicus*, have never been reported in the literature. We hypothesised that *Ae. koreicus* may be an anautogenous mosquito species; however, although autogeny was not present in the studied colony, the phenotype could still be present in different *Ae. koreicus* populations, as previously found for instance in *Ae. albopictus* (Chambers & Klowden, 1994; Mori *et al*., 2008).

The delay of 25.5 hours being observed before mosquito mating could be due to different factors. Although adult female mosquitoes are ready to be inseminated once they emerge, male antennae and genitalia at the moment of imaginal stage emergence are not in the correct morphological conformation to allow copula. Physical changes must occur for the males to become sexually active (Oliva *et al*., 2014a). These changes include the erection of fibrillar hairs in the antennae, (important for female localisation (Roth, 1948)), and the permanent 180° rotation of terminalia part of the genitalia to correctly orient the male genital structure for mating (Lamb, 1922). In particular, the time required for this rotation varies among mosquito species and can take up to four days, for example, as reported in the species *Aedes provocans* (Smith & Gadawski, 1994). The time of *Ae. koreicus* male genitalia rotation is not known, which justifies the choice to cage females with 6-7 days old virgin males. Moreover, mating may be encouraged by behaviours displayed in the wild, such as swarming (Cabrera & Jaffe, 2007), that are challenging to create in a laboratory colony.

In this preliminary exploration of *Ae. koreicus* and *Ae. albopictus* mating interactions, *Ae. albopictus* males showed repeated and aggressive mating attempts towards *Ae. koreicus* but were unable to transfer sperm to *Ae. koreicus*. The different sizes of the two species might be one explanation for how this has played a role in the outcome of this experiment, with the wing length for females of *Ae. koreicus* being reported over 3mm and *Ae. albopictus* male wing length approximately 2mm (Baldacchino *et al*., 2017; Ciocchetta *et al*., 2017; Pudar *et al*., 2021). Yet, the lack of sperm doesn’t necessarily exclude a satyrization effect produced by *Ae. albopictus* males, because the transfer of male accessory glands products (responsible for the satyrization effect) may occur even in the absence of sperm in the spermathecae, as demonstrated by Carrasquilla and Lounibos (Carrasquilla & Lounibos, 2015). Although satyrization between these two species seems unlikely, the aggressive mating attempts shown by *Ae. albopictus* males towards *Ae. koreicus* females could prevent less aggressive *Ae. koreicus* males from mating, and therefore lead to a decrease in *Ae. koreicus* numbers in the field.

Samples tested for *Wolbachia* were collected during the early stages of *Ae. koreicus* invasion in Italy. *Wolbachia* was not detected in *Ae. koreicus* from the first established field population in Belluno from which the studied colony was derived. Therefore, the bacterial endosymbiont is unlikely to have affected *Ae. koreicus* reproductive behaviour in the initial establishment of this mosquito in Italy. The absence of *Wolbachia* in *Ae. koreicus* has been confirmed by subsequent studies in immature and adult stages sampled in the Province of Trento in 2015 and 2017 (Rosso *et al*., 2018; Alfano *et al*., 2019). It should be noted that in our preliminary investigation we tested a small sample (N=21) of *Ae. koreicus* females; nonetheless, our results suggest that even if present in the mosquito population initially established in Italy, the prevalence of *Wolbachia* was low. Interestingly, a more recent study (Damiani *et al*., 2022) on samples collected between 2019 and 2020 in several villages in North-East Italy detected *Wolbachia* in 2 of the 85 females examined. Further research with larger sample sizes could help establish whether *Wolbachia* is present at a very low prevalence in the *Ae. koreicus* population established in Italy or whether the recent discovery is due to introgression (Bargielowski *et al*., 2015) from other mosquito species carrying the endosymbiont (*Wolbachia* strains isolated in *Ae. koreicus* are closely related to the *w*AlbB strain, one of the two native strains of *Ae. albopictus* (McMeniman *et al*., 2009). In the Province of Trento for instance, *Ae. albopictus* had 2.5% prevalence of *Wolbachia* (Rosso *et al*., 2018).

Considering the importance played by reproductive success in ensuring the establishment and growth of invasive mosquito populations during the colonization of new territories, our preliminary results are aimed at informing further studies to assist in determining the invasive potential of *Ae. koreicus* and the public health risk posed in areas of recent introduction.

## Acknowledgements

We thank Dr. Oselyne Ong for her help in screening *Ae. koreicus* samples for the presence of *Wolbachia pipientis* and Dr. Jonathan Darbro for his help in maintaining the mosquito colony.

## Author Contributions

Conceived and designed the study: SC, FDF and GJD. Collected the data: SC. Analysed the data: SC. Drafted the manuscript: SC. Reviewed the manuscript: FDF, FM, GC and GJD. All authors read and approved the final manuscript.

## Data Availability

The data supporting the conclusions of this article are included within the article and its additional files.

## Supporting Information

**Ae. koreicus mating.mp4** mp4 video, 7.92 MB *Ae. koreicus* males and females in the act of copula

**Ae. albopictus_Ae.koreicus interaction.mp4** mp4 video, 3.07 MB Repeated interactions between *Ae. koreicus* female and *Ae. albopictus* males

